# Spike timing in the attention network predicts behavioral outcome prior to target selection

**DOI:** 10.1101/2020.04.03.024109

**Authors:** Ian C. Fiebelkorn, Sabine Kastner

## Abstract

There has been little evidence linking changes in spiking activity that occur prior to a spatially predictable target (i.e., prior to target selection) to behavioral outcomes, despite such preparatory changes being widely assumed to enhance the sensitivity of sensory processing. We simultaneously recorded from frontal and parietal nodes of the attention network, while macaques performed a spatial-cueing task. When anticipating a spatially predictable target, different patterns of coupling between spike timing and oscillatory phase in local field potentials—but not changes in spike rate—were predictive of different behavioral outcomes. These behaviorally relevant differences in local and between-region synchronization occurred among specific cell types that were defined based on their sensory and motor properties, providing insight into the mechanisms underlying enhanced sensory processing prior to target selection. We propose that these changes in neural synchronization reflect differential, anticipatory engagement of the network nodes and functional units that shape attention-related sampling.

## INTRODUCTION

Survival depends on successfully navigating a complex, dynamic environment, but the brain has limited processing resources for the sampling of environmental information (Desimone and Duncan, 1995; Kastner and Ungerleider, 2000). Selective attention comprises a set of neural mechanisms through which some aspects of the environment receive preferential processing relative to others. For example, selective attention can boost sensory processing at a behaviorally relevant location (Helmholtz, 1867; Posner, 1980). This spatially specific boost in sensory processing, termed spatial attention, is characterized by various changes in neural activity, either in response to a target or in anticipation of a target (Buschman and Kastner, 2015; Reynolds and Chelazzi, 2004). Here, we will specifically focus on neural activity that occurs in preparation for a spatially predictable target, in the absence of additional sensory stimulation (i.e., during a variable cue-target delay) and before target selection. Such attention-related, preparatory changes in neural activity occur across multiple levels of observation, from single neurons (Luck et al., 1997) to large-scale neural populations (Kastner et al., 1999).

Preparatory (or ‘pre-target’) changes in neural activity during the deployment of spatial attention are thought to enhance the sensitivity and efficiency of sensory processing (Kastner et al., 1999; Luck et al., 1997), subsequently leading to target selection and attention-related improvements in behavioral performance (Posner, 1980). Consistent with this common interpretation, pre-target neural signals at the population level that derive from summed synaptic potentials (i.e., local field potentials [LFPs] and LFP-like signals) are indeed predictive of behavioral performance (Busch and VanRullen, 2010; Fiebelkorn et al., 2018, 2019; Gonzalez Andino et al., 2005; Hanslmayr et al., 2007; Rohenkohl et al., 2018; Thut et al., 2006). For example, enhanced visual-target detection at an attended location is associated with the synchronization and desynchronization of LFP and LFP-like signals within specific frequency bands (e.g., desynchronization within the alpha band [9–13 Hz] (Thut et al., 2006) and synchronization within the gamma band [35–55 Hz] (Fiebelkorn et al., 2018; Gonzalez Andino et al., 2005)).

In comparison, there is little evidence linking attention-related changes in pre-target spiking activity to behavioral outcomes. While previous studies have repeatedly demonstrated attention-related modulation of pre-target spike rates among neurons in visual cortex (Luck et al., 1997; Reynolds and Chelazzi, 2004), there is seemingly no relationship between such preparatory effects and attention-related improvements in behavioral performance (Galashan et al., 2013; Womelsdorf et al., 2006; Zenon and Krauzlis, 2012). For example, trials with higher pre-target spike rates among neurons representing the attended location are not predictive of better behavioral performance (e.g., faster response times) when targets subsequently occur at the attended location. These attention-related findings contrast with findings associated with other cognitive processes, such as working memory, where spike rates during a memory delay are predictive of behavioral outcomes (Constantinidis et al., 2018).

Attention-related neural effects in visual cortex, such as elevated spike rates during a cue-target delay (i.e., following a spatial cue but prior to target presentation), are generated by feedback signals from a large-scale network of higher-order cortical and subcortical structures (i.e., the attention network) (Corbetta and Shulman, 2002; Kastner and Ungerleider, 2000; Moore and Armstrong, 2003). Attention-related improvements in behavioral performance might therefore also arise from this attention network. For example, pre-target microstimulation in the attention network (i.e., the frontal eye fields) leads to attention-like improvements in behavioral performance (Moore and Fallah, 2001). The specific neural mechanisms leading to these improvements in behavioral performance, however, remain poorly understood. Here, we investigated the link between behavioral outcomes and various aspects of pre-target spiking activity in two well-established cortical nodes of the macaque attention network: the frontal eye fields (FEF) (Fiebelkorn and Kastner, 2020; Squire et al., 2013) and the lateral intraparietal area (LIP) (Bisley and Goldberg, 2010; Fiebelkorn and Kastner, 2020).

Classical views of cortical function attribute computation and information coding to spike rate (Averbeck et al., 2006; Shadlen and Newsome, 1998). Consistent with this viewpoint, an elevated pre-target spike rate in the attention network reflects the maintenance of spatial attention at a specific location (Fiebelkorn et al., 2018, 2019). That is, an elevated pre-target spike rate specifically occurs among neurons with RFs that overlap with the attended location, perhaps maintaining a representation of the to-be-attended location (Luck et al., 1997). Here, we first examined whether differences in pre-target spike rates in FEF and LIP— unlike pre-target spike rates in visual cortex (Galashan et al., 2013; Womelsdorf et al., 2006; Zenon and Krauzlis, 2012)—are associated with differences in behavioral outcomes.

In addition to examining the magnitude of spiking activity (i.e., spike rate) during preparatory attentional deployment, we examined a potential link between the temporal pattern of pre-target spiking activity and behavioral performance. Specifically, we hypothesized that oscillatory patterns in spiking activity might reflect behaviorally relevant, preparatory changes in functional connectivity. FEF and LIP are positioned at a nexus of sensory and motor processes, directing both attention-related boosts in sensory processing and exploratory movements (e.g., saccadic eye movements) (Fiebelkorn and Kastner, 2019). Pre-target changes in functional connectivity (Bosman et al., 2012; Buschman and Miller, 2007; Fiebelkorn et al., 2018, 2019; Gregoriou et al., 2009; Rohenkohl et al., 2018; Saalmann et al., 2007; Saalmann et al., 2012; Voloh et al., 2015) might therefore optimize behavior by priming the attention network in anticipation of a spatially predictable target, leading to enhanced sensory processing and more efficient sensorimotor integration when the target occurs.

## RESULTS

Two monkeys performed a spatial-cueing task (Figure 1A), with a peripheral cue indicating the location where a visual target was most likely to occur (i.e., with 78% cue validity). While we observed significant behavioral effects in both RTs and hit rates (Figure 1B), we focused on the relationship between spiking activity and RTs for subsequent analyses, dividing trials into those resulting in faster or slower RTs. This measure of behavioral performance thus provided an equal number of trials for each condition (i.e., faster RTs vs. slower RTs), allowing for unbiased between-condition comparisons (see Online Methods). We further limited our analyses to spiking activity among visually responsive (i.e. visual and visual-movement) neurons, as previous studies have shown that elevated spiking activity in the attention network during a cue-target delay only occurs among neurons with visual-sensory responses, and not among neurons exhibiting solely motor responses (Fiebelkorn et al., 2019; Gregoriou et al., 2012; Thompson et al., 2005). All of the subsequent analyses focus on spiking activity during the variable cue-target delay, prior to target selection (i.e., pre-target spiking activity).

**Figure 1.**
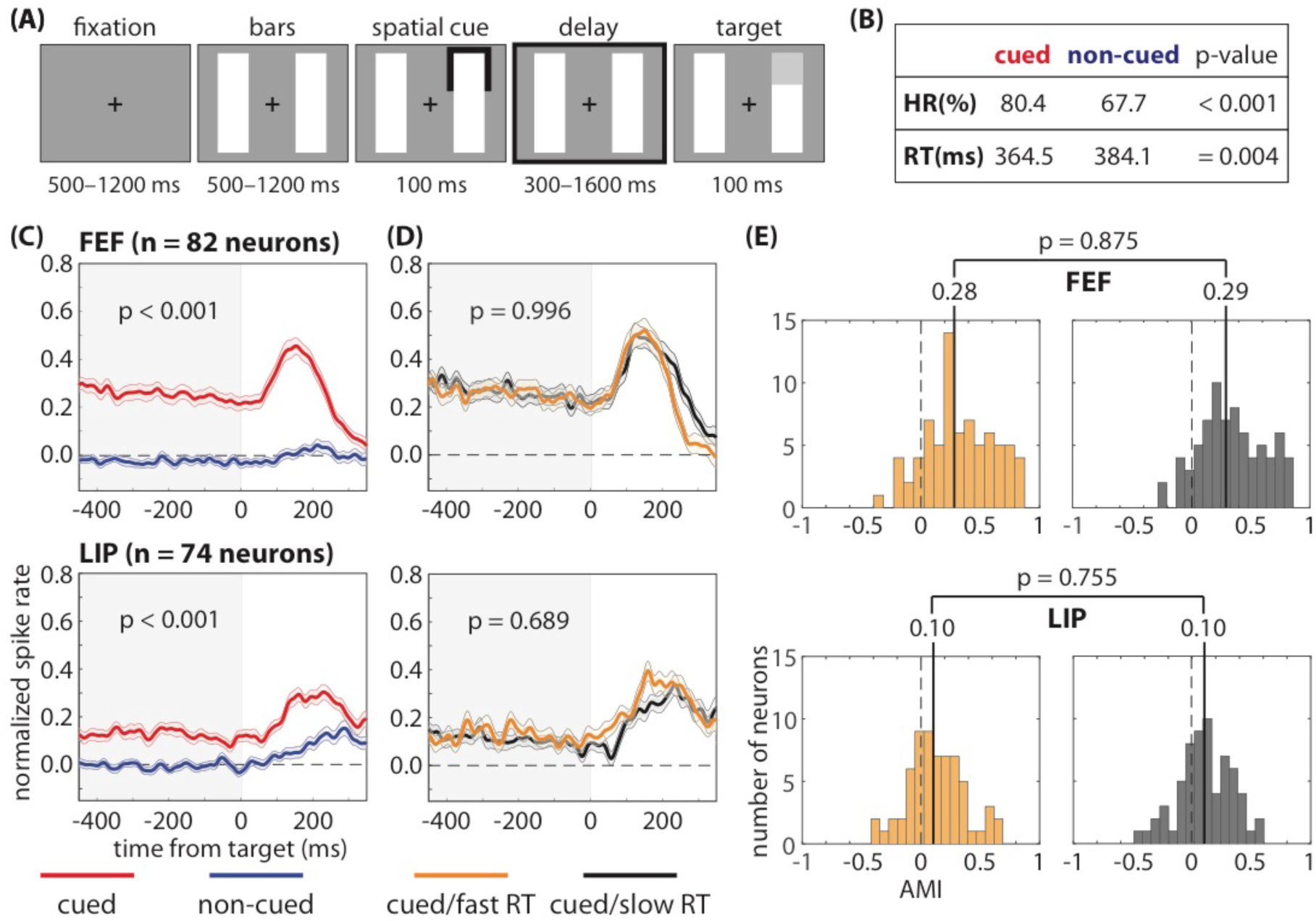
Pre-target differences in spike rates are not associated with differences in response times. We recorded from the frontal eye fields (FEF) and the lateral intraparietal (LIP) region while monkeys completed **(A)** a spatial-cueing task. The animals demonstrated **(B)** both a higher hit rate (HR) and faster response times (RTs) when low-contrast targets occurred at the cued location relative to a non-cued location. **(C)** Pre-target spike rates (i.e., during the cue-target delay), averaged across neurons, were higher when receptive fields (RFs) overlapped the cued location; however, **(D)** we observed no differences in pre-target spike rates, averaged across neurons, between fast- and slow-RT trials (i.e., when RFs overlapped the cued location). **(E)** We also observed no differences in the median attentional modulation index (AMI) between fast- and slow-RT trials. The shaded area around each line represents the standard error of the mean.

### Pre-target spike rates are not associated with behavioral performance

Elevated spike rates in response to sensory stimulation (i.e., in response to a target) increase the signal-to-noise ratio, improving signal detection and speeding RTs (Galashan et al., 2013; Womelsdorf et al., 2006). The functional role of elevated spike rates prior to target selection (i.e., in anticipation of a spatially predictable target and in absence of sensory input) is less clear, as is the relationship between pre-target spike rates and subsequent behavioral performance. Here, we investigated this relationship in two well-characterized nodes of the attention network in macaques: FEF (Fiebelkorn and Kastner, 2020; Squire et al., 2013) and LIP (Bisley and Goldberg, 2010; Fiebelkorn and Kastner, 2020). As a first step, we confirmed a significant increase in spike rates following the cue and prior to target presentation (i.e., during the cue-target delay), specifically when RFs overlapped the cued location relative to when RFs overlapped a non-cued location (Wilcoxon rank-sum test, p < 0.001 for FEF and p < 0.001 for LIP; Figure 1C). We then split trials when RFs overlapped the cued location into two bins based on the median RT (Figure 1D) (Womelsdorf et al., 2006). Figure 1 shows nearly identical mean spike rates (N = 82 for FEF and N = 74 for LIP) for trials that resulted in either faster or slower RTs (Wilcoxon rank-sum test, p = 0.996 for FEF and p = 0689 for LIP). See Supplemental Figure S1 for similar results when binning trials based on whether they resulted in either a hit or a miss.

We also calculated an attentional modulation index for each neuron (see Online Methods). Figure 1E shows similar distributions for trials that resulted in either faster or slower RTs (Wilcoxon rank-sum test, p = 0.875 for FEF and p = 0.755 for LIP), with nearly identical median values in both FEF (i.e., 0.28 and 0.29 for fast- and slow-RT trials, respectively) and LIP (i.e., 0.10 and 0.10 for fast- and slow-RT trials, respectively). The present results therefore suggest that pre-target spike rates in FEF and LIP are not a good predictor of behavioral outcomes. These findings are consistent with previous findings in visual cortex, which similarly found no apparent relationship between trial-to-trial differences in pre-target spike rates and behavioral outcomes (Galashan et al., 2013; Womelsdorf et al., 2006; Zenon and Krauzlis, 2012).

We further explored the relationship between pre-target spike rates and behavioral performance on both a smaller scale (i.e., cell-type specific spiking) and a larger scale (i.e., population spiking). First, we split visually responsive neurons into two functionally defined cell types: neurons that only responded to visual stimulation (i.e., visual neurons) and neurons that both responded to visual stimulation and demonstrated saccade-related activity (i.e., visual-movement neurons) (Fiebelkorn et al., 2018). Here, we tested whether the lack of an apparent relationship between pre-target spike rates and behavioral performance (Figure 1D, E) resulted from combining cell types that are functionally different. Splitting visually responsive neurons into visual and visual-movement neurons, however, similarly revealed no significant relationship between pre-target spike rates and RTs (Figure 2).

**Figure 2.**
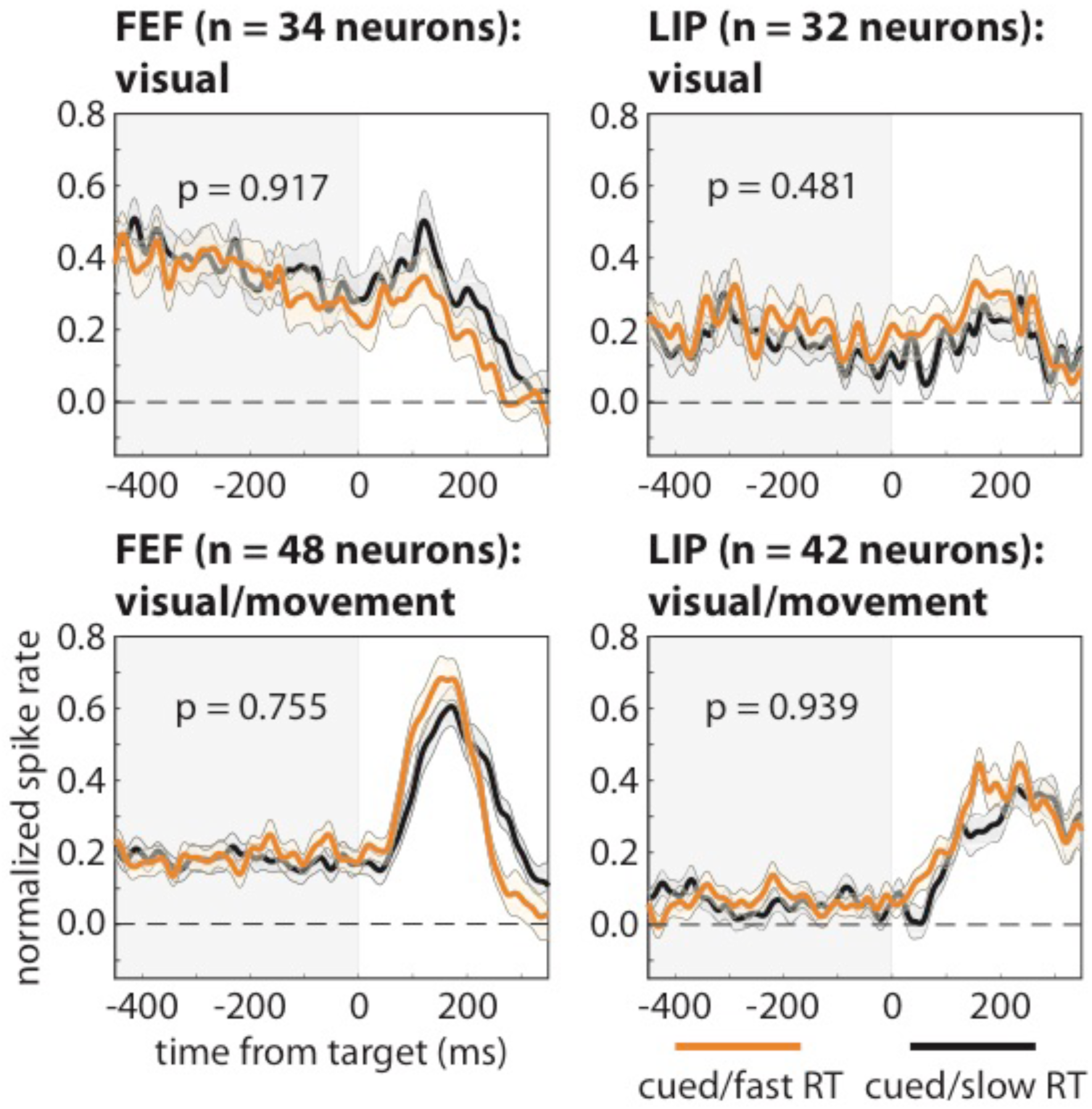
Pre-target differences in spike rate among functionally defined cell types are not associated with differences in response times. We binned neurons in FEF and LIP into visual and visual-movement cell types, with the former only responding to visual stimulation and the latter both responding to visual stimulation and demonstrating saccade-related activity. Regardless of cell type and brain region, we observed no differences in pre-target spike rates, averaged across neurons, between trials that resulted in either faster (orange lines) or slower (black lines) response times (i.e., when RFs overlapped the cued location). The shaded area around each line represents the standard error of the mean.

Next, we measured high-frequency band (HFB) activity in the LFPs (70–150 Hz) as a proxy for population spiking during each recording session (see Online Methods) (Ray and Maunsell, 2011). Although there was no apparent relationship between spike rates and behavioral outcomes at the level of single neurons (Figures 1 and 2), such a relationship might emerge when measuring population-spiking activity. Previous studies have linked lower-frequency population measures in LFPs and LFP-like signals (e.g., EEG) to behavioral performance (Busch and VanRullen, 2010; Fiebelkorn et al., 2018, 2019; Gonzalez Andino et al., 2005; Hanslmayr et al., 2007; Rohenkohl et al., 2018; Thut et al., 2006), but the present results, based on HFB activity (Figure 3), mirrored those based on single-unit activity (Figures 1 and 2). That is, we detected significant attention-related differences (Fig. 3A; Wilcoxon rank-sum test, p < 0.001 for FEF and p = 0.008 for LIP) but no differences based on whether trials at the cued location resulted in either faster or slower RTs (Fig. 3B; Wilcoxon rank-sum test, p = 0.533 for FEF and p = 0.909 for LIP). See Supplemental Figure 2 for similar results when comparing trials that resulted in either a hit or a miss (Wilcoxon rank-sum test, p = 0.322 for FEF and p = 0.531 for LIP). These findings therefore provide further evidence that the lack of a relationship between pre-target spike rates and subsequent RTs (Figure 1D, E) is not an issue of the measurement scale (Cohen and Maunsell, 2010). In summary, we did not find evidence of a relationship between either (i) RTs and spike rates among functionally defined cell types (i.e., visual neurons and visual-movement neurons; Figure 2B) or (ii) RTs and the magnitude of population spiking (Figure 3C).

**Figure 3.**
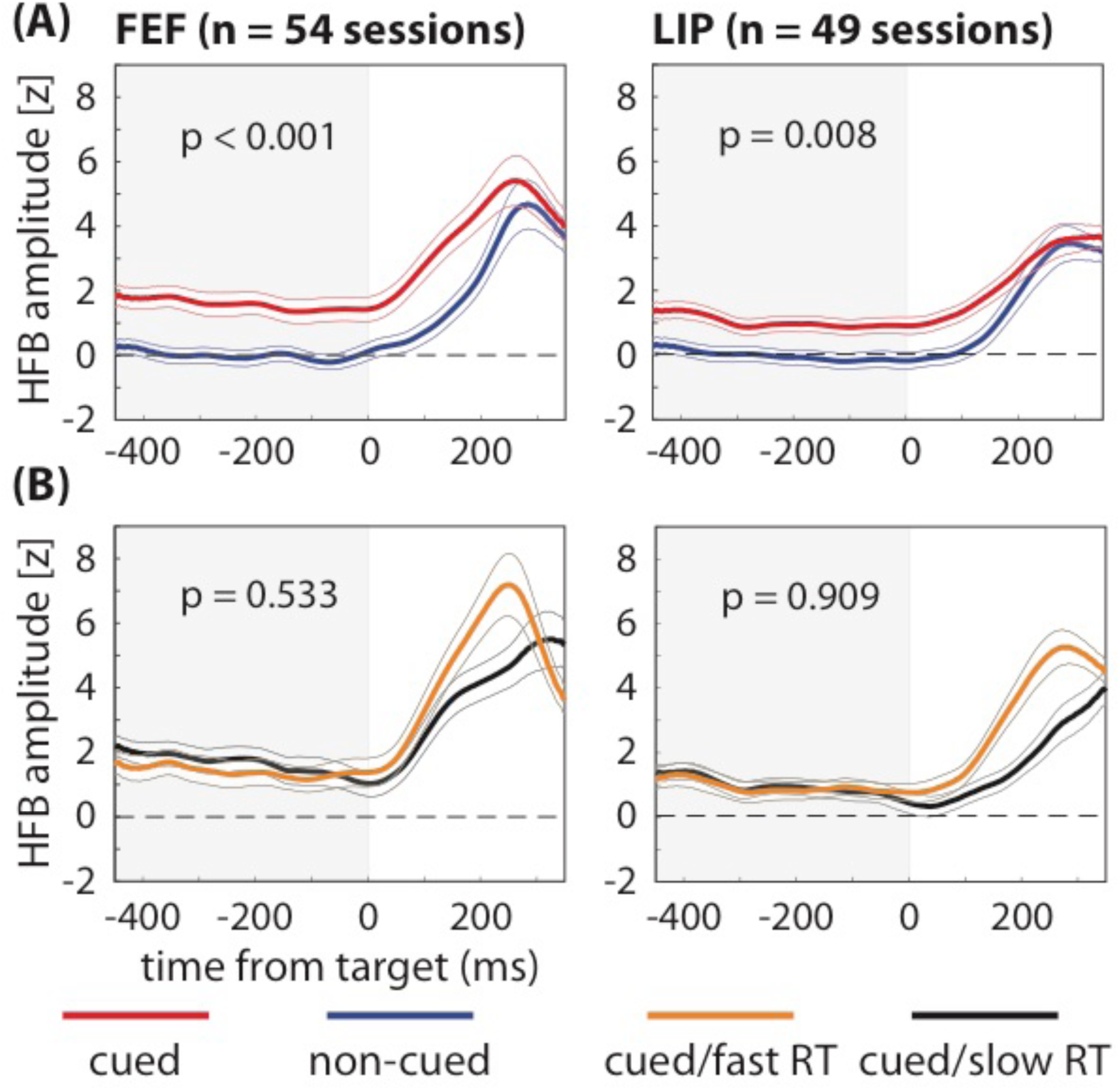
Differences in pre-target population spiking are not associated with differences in response times. **(A)** While we observed significantly greater high-frequency band (HFB) activity—a proxy for population spiking—during the cue-target delay when response fields (i.e., the LFP equivalent of receptive fields) overlapped the cued location (red lines) relative to when response fields overlapped the non-cued location (blue lines), **(B)** we observed no differences between trials that resulted in either faster (orange lines) or slower (black lines) response times (RTs). That is, we observed significant attention-related increases in HFB activity, but no differences associated with behavioral performance (fast-RT versus slow-RT trials). The shaded area around each line represents the standard error of the mean.

### Oscillatory patterns in pre-target spiking activity are associated with behavioral performance

Given the lack of evidence that spike rates prior to target selection influence behavioral performance thus far observed in our data, we examined potential links between other aspects of pre-target spiking activity and behavioral performance. We first used the Fano factor to examine whether spike-count variability influences RTs (Supplemental Figure 3). The Fano factor measures the consistency of spike counts across trials, with decreased response variability (i.e., more consistent spike counts) potentially improving the coding of sensory information in neural signals (Mitchell et al., 2009). We did not, however, find a significant difference in Fano factor between fast- and slow-RT trials when RFs overlapped the cued location (Wilcoxon rank-sum test, p = 0.959 for FEF and p = 0.766 for LIP; see also Chang et al. (2012); (Mitchell et al., 2009)). Similar to pre-target spike rates, the present results indicate that pre-target spike-count variability (i.e., during the cue-target delay) is not associated with subsequent behavioral performance.

We next examined whether spike timing influences RTs, focusing on the relationship between oscillatory synchronization in pre-target spiking activity and behavioral performance. Such synchronization reflects the temporal coordination of local and between-region neural activity (Buschman and Kastner, 2015), and synchronization at various frequencies has been repeatedly associated with specific functions (Bastos et al., 2015; Fiebelkorn and Kastner, 2019; Fries, 2009; Jensen and Mazaheri, 2010). For example, increased synchronization in the gamma band (> 35 Hz) has been linked to enhanced sensory processing (Fries, 2009), while synchronization in the alpha band (9–14 Hz) has been linked to diminished sensory processing (Foxe and Snyder, 2011). Because the spiking of single neurons can be sparse on a given trial, oscillatory patterns in spiking activity are typically assessed by measuring whether spikes cluster at specific phases of oscillatory activity in the LFPs. This relationship between neuronal spikes and oscillatory phase in LFPs is referred to as spike-LFP phase coupling and is an indication of spike timing control.

Figure 4A compares local spike-LFP phase coupling between trials when RFs overlapped either the cued or a non-cued location. Similar to previous studies (Buschman and Miller, 2007; Fiebelkorn et al., 2018), we found that the deployment of spatial attention was associated with both increased beta-band (26–36 Hz) synchronization in FEF and increased gamma-band (44–54 Hz) synchronization in LIP (permutation test, p < 0.05, corrected for multiple comparisons). Figure 4B compares local spike-LFP phase coupling between trials that resulted in either faster or slower RTs when RFs overlapped the cued location (i.e., under conditions of attentional deployment). Unlike the previous analyses, which focused on spike rates or spike-count variability (Figures 1–3, Supplemental Figure 3), different oscillatory patterns in spiking activity were associated with different behavioral outcomes. That is, fast-RT trials were characterized by significantly higher beta synchronization (i.e., 19–26 Hz) in FEF and significantly lower alpha synchronization (i.e., 9–15 Hz) in LIP (permutation test, p < 0.05, corrected for multiple comparisons). As shown in Figure 4A, attentional deployment at the cued location was associated with an increase in gamma synchronization in LIP; however, Figure 4B shows that there was no significant difference in gamma synchronization between fast- and slow-RT trials. Supplemental Figure 4 demonstrates similar results when measuring oscillatory patterns in the HFB signal (as a proxy for population spiking).

**Figure 4.**
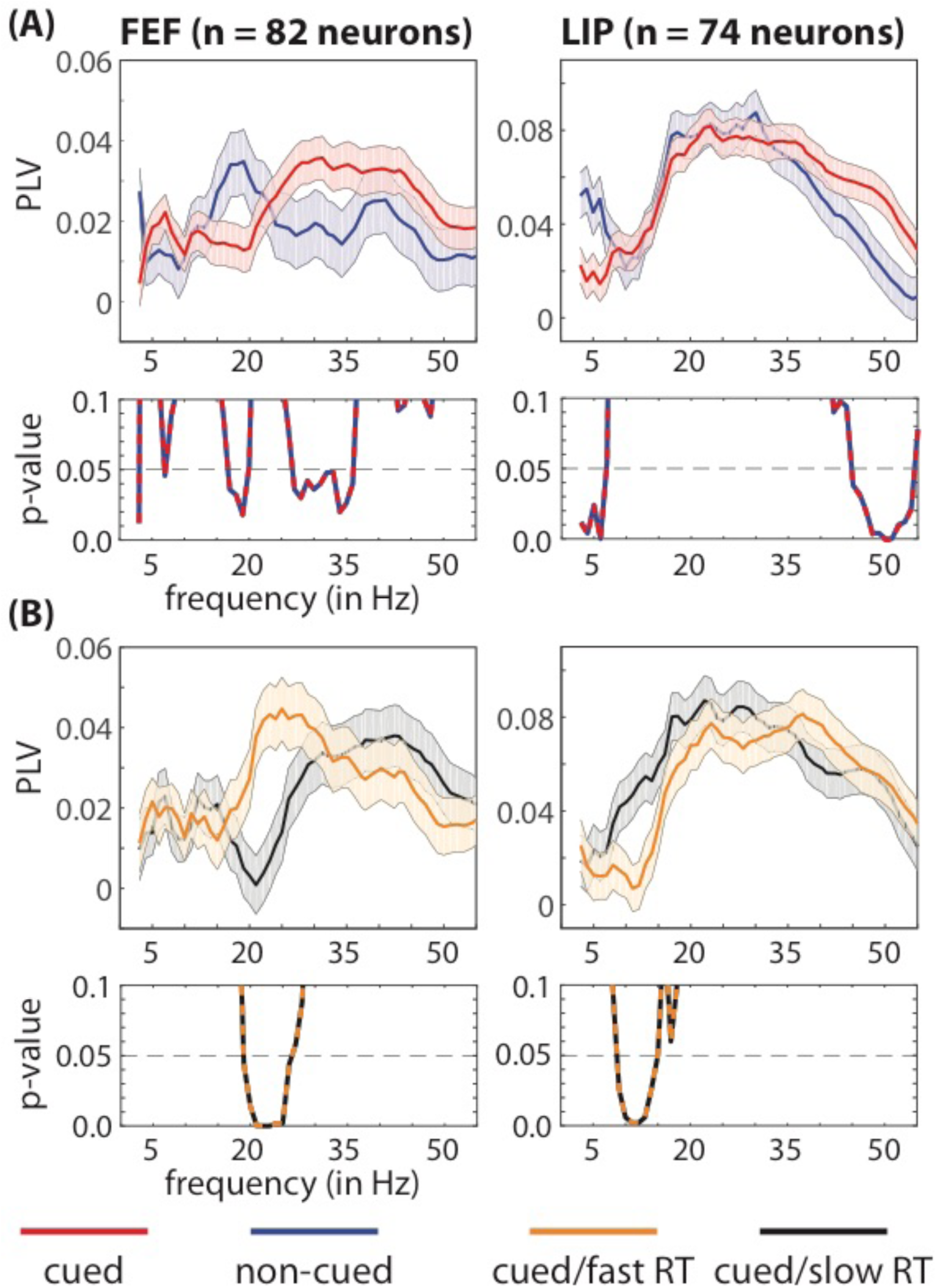
Faster response times are characterized by specific oscillatory patterns in pre-target spiking activity, locally in FEF and LIP. **(A)** We observed significant local differences in the frequencies associated with clustering between spikes and oscillatory phase in LFPs (i.e., spike-LFP phase coupling), depending on whether receptive fields and response fields overlapped either a cued (red lines) or a non-cued (blue lines) location. **(B)** We also observed significant differences between trials that resulted in either faster (orange lines) or slower (black lines) response times (i.e., when receptive fields and response fields overlapped the cued location). The p-values for between-condition comparisons are represented below each panel. The shaded area around each line represents the standard error of the mean.

To examine whether behaviorally relevant differences in oscillatory synchronization were associated with specific functions, we next measured local spike-LFP phase coupling separately for visual and visual-movement neurons. Figure 5 shows that both the increased beta synchronization in FEF and the decreased alpha synchronization in LIP—associated with fast-RT trials—occurred exclusively among visual-movement neurons (permutation test, p < 0.05, corrected for multiple comparisons).

**Figure 5.**
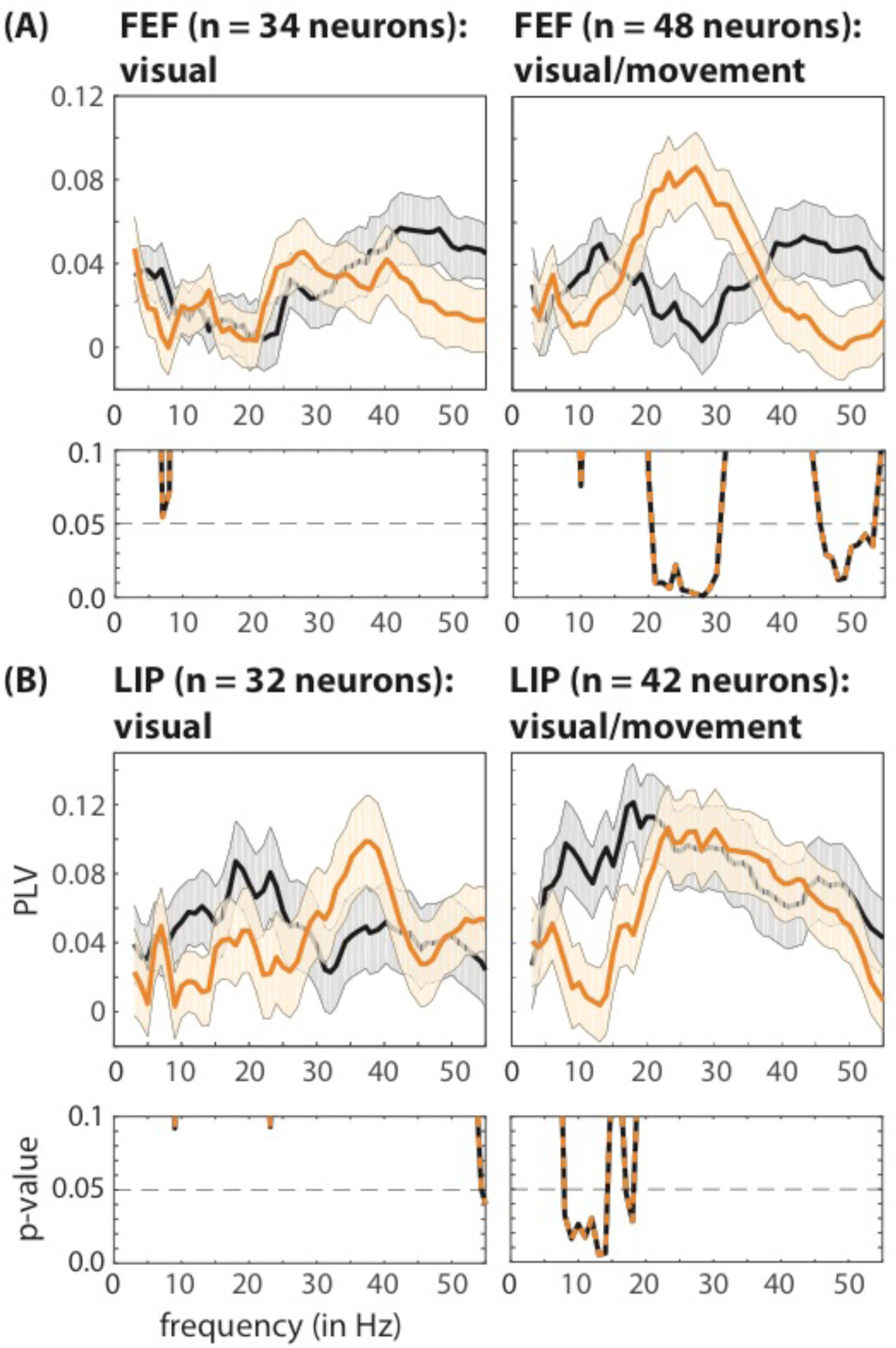
Faster response times are characterized by cell-type specific oscillatory patterns in pre-target spiking activity, locally in FEF and LIP. In both **(A)** FEF and **(B)** LIP, significant differences in local spike-LFP phase coupling between fast- (orange lines) and slow-RT (black lines) trials occurred exclusively among visual-movement neurons (i.e., neurons with both visual-sensory responses and saccade-related activity). The p-values for between-condition comparisons are represented below each panel. The shaded area around each line represents the standard error of the mean.

We next examined between-region synchronization between cell-type specific spiking activity and oscillatory phase in the LFPs. The synchronization of spiking activity in one brain region and frequency-specific oscillatory phase in another brain region is often interpreted as evidence of network-level participation in a common function. Here, we examined whether such between-region functional connectivity differed between trials that resulted in either faster or slower RTs (when RFs overlapped the cued location). We again measured spike-LFP phase coupling separately for visual and visual-movement neurons (Figure 6). Fast-RT trials were characterized by significantly higher synchronization between spikes in FEF and both theta- (3–7 Hz) and beta-band (19 and 22–27 Hz) activity in LIP (permutation test, p < 0.05, corrected for multiple comparisons), with theta synchronization occurring among visual neurons and beta synchronization occurring among visual-movement neurons (Figure 6A). We describe the potential functional significance of these cell-type specific effects in the Discussion section.

**Figure 6.**
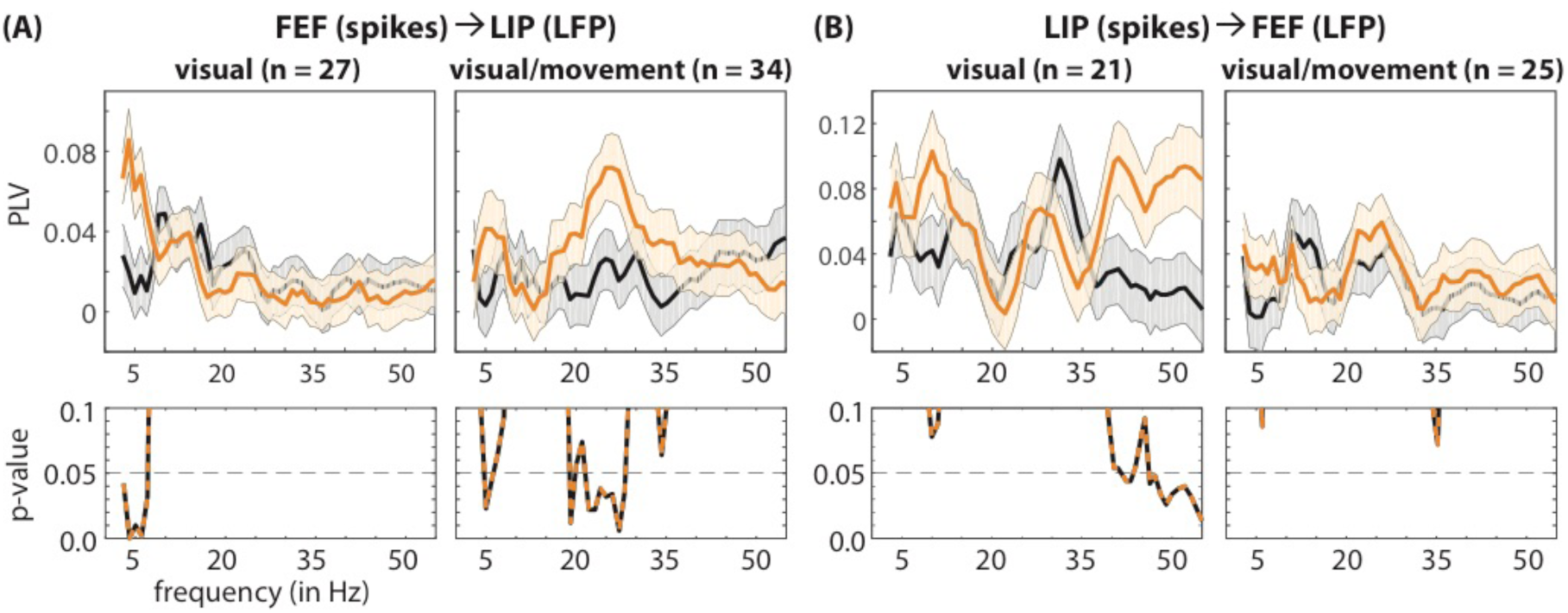
Faster response times are characterized by network-level oscillatory patterns in pre-target spiking activity. **(A)** Fast-RT trials (orange lines) are associated with (i) greater theta synchronization between visual neurons in FEF and LFPs in LIP and (ii) greater beta synchronization between visual-movement neurons in FEF and LFPs in LIP (relative to slow-RT trials, black lines). **(B)** Fast-RT trials are also associated with greater gamma synchronization between visual neurons in LIP and LFPs in FEF. The p-values for between-condition comparisons are represented below each panel. The shaded area around each line represents the standard error of the mean.

Unlike within-region gamma synchronization (Figure 4B), between-region gamma synchronization was associated with differences in behavioral performance (Figure 6B). That is, we previously noted that attentional deployment was associated with increased gamma synchronization locally in LIP; however, there was no difference in gamma synchronization between fast- and slow-RT trials (Figure 4B). In comparison, at the network level, fast-RT trials (relative to slow-RT trials) were characterized by significantly higher synchronization between spikes in LIP and gamma-band (38 and 47–55 Hz) activity in FEF (permutation test, p < 0.05, corrected for multiple comparisons), occurring exclusively among visual neurons (i.e., neurons with a visual-sensory response but no saccade-related activity). These results are therefore consistent with previous work linking gamma synchronization to attention-related boosts in visual-sensory processing (Gregoriou et al., 2009; Womelsdorf et al., 2006).

Because differences in oscillatory power in LFPs might spuriously lead to differences in spike-LFP phase coupling (see Online Methods), we also conducted a control analysis, equating oscillatory power between fast- and slow-RT trials. Those results, illustrated in Supplemental Figure 5, confirmed our findings, revealing significantly different oscillatory synchronization in cell-type specific spiking activity associated with either faster or slower RTs, at both the local and the network levels.

### Links between oscillatory patterns in pre-target spiking activity and behavioral performance reflect alternating attentional states

The present results indicate that specific oscillatory patterns in pre-target spiking activity are associated with either faster or slower RTs. These oscillatory patterns are associated with specific, functionally defined cell types and occur across multiple frequency bands. We previously demonstrated that these same frequency bands define two rhythmically alternating attentional states associated with either better or worse visual-target detection at a spatially cued location (Fiebelkorn et al., 2018).

Whereas classic views assumed that spatial attention samples the visual environment continuously, recent research has instead provided evidence that spatial attention samples the visual environment in rhythmic cycles (Landau and Fries, 2012; Pringsheim et al., 2013; VanRullen et al., 2007). Such rhythmic sampling is associated with theta-band activity in the attention network (Fiebelkorn and Kastner, 2019; Fiebelkorn et al., 2018, 2019; Helfrich et al., 2018), with different theta phases in the LFPs linked to alternating periods of either better or worse behavioral performance. Theta-dependent periods associated with better visual-target detection at the cued location (i.e., during the ‘good’ theta phase) are characterized by increased beta- and gamma-band activity in FEF and LIP, while theta-dependent periods associated with worse visual-target detection at the cued location (i.e., during the ‘poor’ theta phase) are characterized by increased alpha-band activity in LIP (Fiebelkorn and Kastner, 2019; Fiebelkorn et al., 2018).

Based on previous findings that higher LFP frequencies (i.e., alpha-, beta-, and gamma-band activity) in the attention network are temporally organized by theta-band activity (Fiebelkorn and Kastner, 2019; Fiebelkorn et al., 2018, 2019; Helfrich et al., 2018), we hypothesized that behaviorally relevant oscillatory synchronization in spiking activity should be similarly tied to the phase of theta-band activity. Figure 7 (A, B) shows spike-LFP phase coupling in FEF and LIP as a function of time, isolating the frequency bands where we detected statistically significant differences between fast- and slow-RT trials (Figure 4B). These results indicate that (i) pre-target beta (19–26 Hz) synchronization in FEF was generally stronger (i.e., at different pre-target time points) on fast-RT trials (Figure 7A) and (ii) that pre-target alpha (9–15 Hz) synchronization in LIP was generally stronger on slow-RT trials (Figure 7B). A closer look, however, further suggests that the strength of spike-LFP phase coupling waxed and waned during the cue-target delay, seemingly at a theta rhythm (3–8 Hz). We therefore tested whether spike-LFP phase coupling at these higher frequencies (i.e., beta and alpha) was linked to the phase of theta-band activity (Figure 7C, D). Here, we calculated spike-LFP phase coupling (i.e., at beta and alpha) as a function of theta phase. We then fit the resulting functions (i.e., spike-LFP phase coupling as a function of theta phase) with a one-cycle sine wave, using the amplitude of that sine wave to measure the strength of the relationship between theta phase and spike-LFP phase coupling at higher frequencies (see Online Methods) (Fiebelkorn et al., 2019). While the relationship between theta phase and alpha synchronization in LIP was not statistically significant (permutation test, p = 0.13, corrected for multiple comparisons), there was a statistically significant relationship between theta phase and beta synchronization in FEF (permutation test, p < 0.05, corrected for multiple comparisons). The results in FEF are therefore consistent with theta-dependent, alternating attentional states associated with either faster or slower RTs: with faster RTs tending to occur when a low-contrast visual-target is presented during the theta-dependent attentional state characterized by higher beta synchronization in FEF, and slower RTs tending to occur when a low-contrast visual-target is presented during the attentional state associated with lower beta synchronization in FEF.

**Figure 7.**
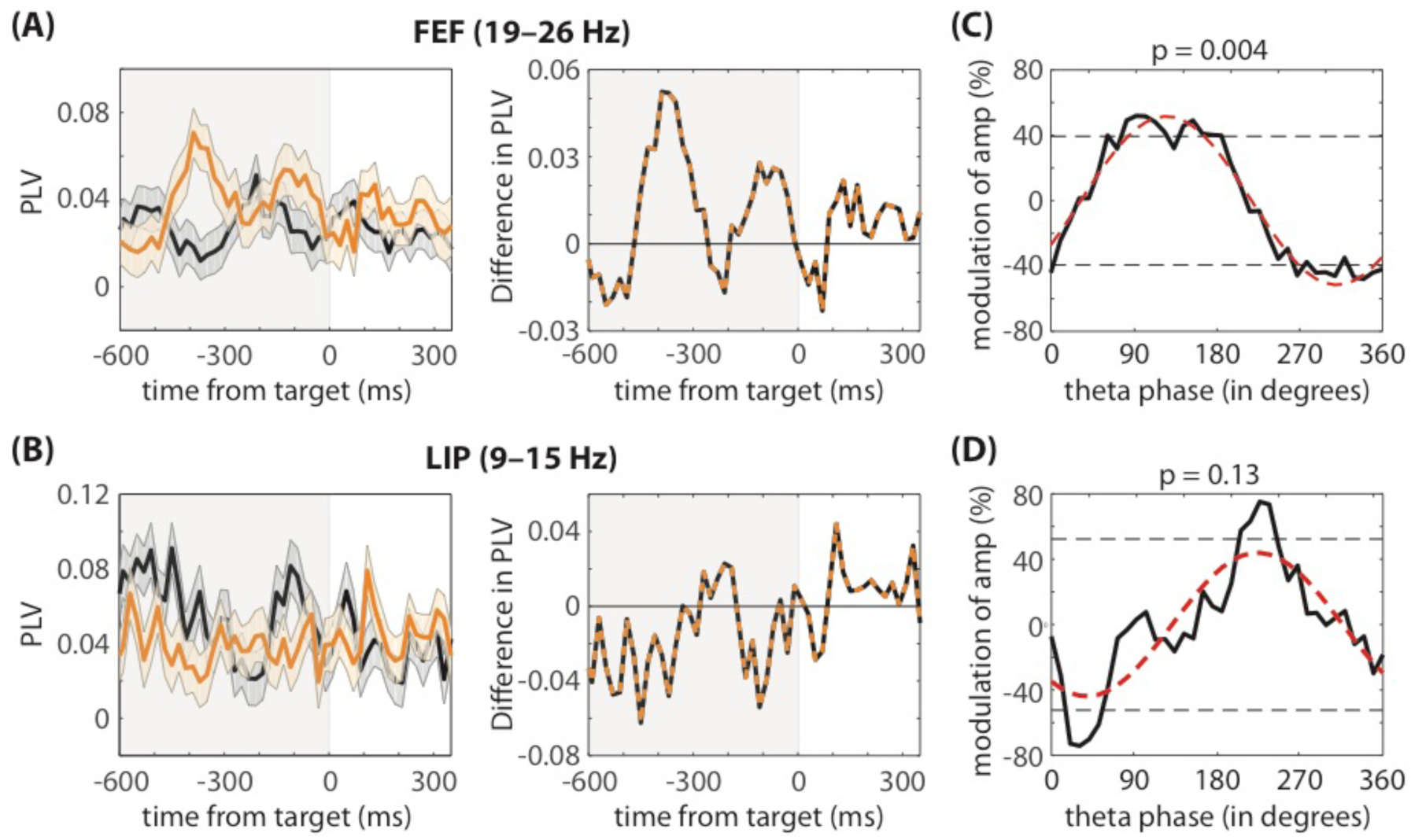
Functional connectivity is highly dynamic during attentional deployment. Although **(A)** beta synchronization in FEF is generally higher on fast-RT trials (orange lines) and **(B)** alpha synchronization in LIP is generally higher on slow-RT trials (black lines); however, the difference between fast- and slow-RT trials (orange and black lines) reveals a pattern consistent with previous studies linking theta-band activity in the attention network to fluctuations in behavioral performance. **(C)** In FEF, the phase of theta-band activity (at 4Hz) significantly modulated spike-LFP phase coupling in the beta band. In LIP, the relationship between the phase of theta-band activity and spike-LFP phase coupling in the alpha band was not statistically significant.

## DISCUSSION

We investigated the link between spiking activity in higher-order nodes of the attention network (i.e., FEF and LIP) and behavioral outcomes during a spatial-cueing task. We specifically examined the behavioral relevance of spiking activity that occurs during preparatory attentional deployment (i.e., during a cue-target delay). Whereas numerous studies have described attention-related changes in target responses to behavioral performance (Buschman and Kastner, 2015), there has previously been little evidence linking behavioral performance to attention-related changes in spiking activity that occur prior to target selection. Spike rate during this time period encodes the focus of spatial attention, with a higher spike rate occurring when RFs overlap the cued location (Figure 1). Consistent with previous recordings in visual cortex (Galashan et al., 2013; Womelsdorf et al., 2006; Zenon and Krauzlis, 2012), however, pre-target spike rates in FEF and LIP were not associated with subsequent behavioral performance (Figures 1–3). In comparison, pre-target differences in spike-LFP phase coupling (i.e., the timing of spiking activity relative to oscillatory phase in LPFs) were both indicative of the focus of spatial attention (i.e., spikes were coupled to different frequencies depending on the focus of attention; Figure 4) and associated with subsequent behavioral performance (Figures 4–6).

Spike rate has long been considered a critical component of information encoding in the brain (Ferster and Spruston, 1995; Shadlen and Newsome, 1998), and in contrast to the present findings, previous studies have demonstrated that spike rate is associated with behavioral outcomes during other cognitive processes, such as visual-working memory (Constantinidis et al., 2018). For example, a higher spike rate during a memory delay is associated with correct trials, while a lower spike rate is associated with error trials (Funahashi et al., 1989; Fuster, 1973). The behavioral relevance of pre-target spike rate therefore seems to depend on the cognitive processes engaged by the task at hand (e.g., spatial attention or working memory).

Unlike attention-related changes in pre-target spike rates, the present results demonstrate that local and between-region patterns of pre-target spike-LFP phase coupling are predictive of behavioral outcomes. Spike-LFP phase coupling is an indication of spike-timing control and represents a possible mechanism for temporally organizing neural activity to optimize (i) local computations and (ii) the transfer of information between brain regions (Fiebelkorn and Kastner, 2019; Lisman and Idiart, 1995; Shahidi et al., 2019; Siegel et al., 2009; Voloh et al., 2019). That is, (i) the timing of spiking activity relative to oscillatory phase in LFPs (i.e., spike-LFP phase coupling) can encode information (Kayser et al., 2009; Lisman, 2005; O’Keefe and Recce, 1993; Optican and Richmond, 1987), and (ii) the temporal synchronization of presynaptic neurons can maximize their influence on postsynaptic neurons by allowing for the summation of postsynaptic potentials (Azouz and Gray, 2003; Fries, 2015; Singer and Gray, 1995; Womelsdorf et al., 2007; Zandvakili and Kohn, 2015). Zandvakili and Kohn (2015), for example, reported that spiking activity in the input layers of visual area 2 (i.e., V2) was preceded by a transient increase in coordinated activity in visual area 1 (i.e., V1). Precise spike-timing control might be particularly important for such between-region interactions, where multiple presynaptic neurons target a single postsynaptic neuron. Between-region spike-LFP phase coupling measures the consistency of spike timing in one brain region relative to oscillatory phase in the LFPs (i.e., the frequency-specific synchronization of postsynaptic potentials) of a second brain region, and these between-region interactions are often interpreted as evidence of participation in a common function (i.e., as functional connectivity).

FEF and LIP are positioned at a nexus of the sensory and motor systems, directing both covert (i.e., in the absence of eye movements) and overt (i.e., with eye movements) aspects of environmental sampling (Fiebelkorn and Kastner, 2019, 2020). Here, we propose that the synchronization of pre-target spiking activity at specific oscillatory frequencies in LFPs reflects the engagement of the sensory and motor functions that shape attention-related sampling (Fiebelkorn and Kastner, 2019) and the recruitment of the network nodes that contribute to those functions (Fiebelkorn and Kastner, 2020). For example, increased beta-band activity among visual-movement neurons in FEF has previously been associated with the suppression of attentional shifts (Fiebelkorn et al., 2018; Gregoriou et al., 2012). Consistent with these previous observations, we observed behaviorally relevant beta synchronization at both the local and the network levels exclusively among visual-movement neurons (Figures 5 and 6). That is, higher beta synchronization among visual-movement neurons (rather than visual neurons) was associated with faster RTs. Increased beta synchronization in FEF prior to target presentation (Figure 4) might therefore reflect a suppression of attentional shifts that co-occurs with greater attentional focus at the cued location.

In comparison, we observed behaviorally relevant gamma synchronization, occurring at the network level (i.e., between LIP and FEF), exclusively among visual neurons (Figure 6). Increased gamma synchronization has been repeatedly linked to enhanced processing in feedforward sensory channels (Bastos et al., 2015; Fries, 2015; Womelsdorf et al., 2006). Greater gamma synchronization during the cue-target delay might therefore reflect an opening of feedforward sensory channels that facilitates the subsequent propagation of target-related information. Womelsdorf et al. (2006) previously linked local pre-target gamma synchronization (40–70 Hz) in visual cortex (area V4) with faster RTs. Here, we observed a local attention-related increase in pre-target gamma synchronization, occurring in LIP (Figure 4A). However, unlike gamma synchronization in visual cortex (Womelsdorf et al., 2006), this local gamma synchronization in LIP was not associated with differences in RTs (Figure 4B). Faster RTs were instead only associated with increased gamma synchronization between frontal (spikes) and parietal (LFPs) nodes of the attention network (Figure 6B), highlighting the importance of network-level interactions in determining behavioral outcomes.

In addition to between-region gamma synchronization, faster RTs were associated with between-region theta synchronization that was also exclusively linked to visual neurons in FEF (Figure 6A). Previous research has demonstrated that spatial attention samples the visual environment in theta-rhythmic cycles (4–6 Hz) (Fiebelkorn et al., 2018, 2019; Fiebelkorn et al., 2013; Helfrich et al., 2018; Landau and Fries, 2012; Landau et al., 2015). This theta-rhythmic sampling is characterized by alternating periods of either enhanced or diminished perceptual sensitivity (Fiebelkorn et al., 2018; Fiebelkorn et al., 2013; Landau and Fries, 2012; VanRullen et al., 2007), with fluctuations in perceptual sensitivity being associated with fluctuations in spatiotemporal patterns of functional connectivity across the attention network (Fiebelkorn et al., 2018, 2019). For example, we previously demonstrated that theta-dependent periods of enhanced perceptual sensitivity (i.e., the “good” theta phase) at a cued location are associated with increased beta-band activity in LFPs (Fiebelkorn et al., 2018). The present findings demonstrate that faster RTs—during the same spatial-cueing task (Figure 1A)—are also associated with increased beta synchronization in pre-target spike times (Figures 4 and 6), with beta synchronization being similarly modulated by the phase of theta-band activity (Figure 7). We have previously proposed that theta rhythms in the attention network are organizing neural activity into alternating attentional states: an attentional state associated with sensory functions of the attention network (or sampling) and an attentional state associated with motor functions of the attention network (or shifting) (Fiebelkorn and Kastner, 2019). The present findings provide evidence that spatiotemporal patterns in spiking activity also change in a theta-dependent manner, reflecting these alternating attentional states (see Fiebelkorn et al. (2019) for further evidence).

It is important to note that the specific cell-types and oscillatory patterns that are linked to behavioral outcomes might depend on the behavioral measure. Here, we described cell-type specific patterns of functional connectivity associated with faster or slower RTs. Attention-related changes in perceptual sensitivity, however, might rely on different patterns of functional connectivity across frontal and parietal regions, associated with different functional cell types (e.g. visual neurons rather than visual-movement neurons).

We measured oscillatory patterns in the spiking of single neurons by examining the timing of spikes relative to oscillatory phase in LFPs (i.e., spike-LFP phase coupling). Previous studies have examined the relationship between pre-target spiking activity and behavioral performance by focusing on correlations either between pairs of neurons or among small populations of neurons (Cohen and Maunsell, 2009, 2010; Shahidi et al., 2019). Those studies have often relied on measures that correlate trial-by-trial fluctuations in neuronal responses (e.g., spike-count or ‘noise’ correlations), examining overall spike counts (or rates) during the time period of interest (e.g., a cue-target delay) (Cohen and Kohn, 2011). Such approaches, however, are blind to behaviorally relevant temporal dynamics, which change on a moment-to-moment timescale (Fiebelkorn and Kastner, 2020). In contrast, Ben Hadj Hassen et al. (2019) recently used multi-unit activity to demonstrate that noise correlations in frontal cortex rhythmically fluctuate over time, with periods of lower noise correlations associated with a higher response probability. These findings, like the present findings, are consistent with an emerging view of attention-related sampling as a highly dynamic, fundamentally rhythmic process (Fiebelkorn and Kastner, 2020). The present findings further indicate that the link between population-level measures of attention-related neural signals and behavioral outcomes stems, at least in part, from the temporal, cell-type specific synchronization of local and between-region spiking activity.

Spiking activity that occurs in response to an attended target (i.e., during target selection) is predictive of behavioral outcomes (Buschman and Kastner, 2015). The present results show that changes in spiking activity that occur in preparation for a spatially predictable target are also predictive of behavioral outcomes. While it has been widely assumed that attention-related, preparatory changes in spiking activity (e.g. following a spatial cue and prior to target selection) improve the sensitivity and efficiency of sensory processing, there has been little evidence that such preparatory changes are linked to subsequent behavioral performance. The present results specifically reveal that different oscillatory patterns in pre-target spike times are linked to either faster or slower RTs. While attention-related changes in pre-target spike rates encode information about the focus of spatial attention, behavioral performance seems to depend more on the temporal synchronization of both local and between-region spiking activity. The present findings thus provide insight into the specific mechanisms that enhance the sensitivity and efficiency of sensory processing in preparation for a spatially predictable target. The sensory and motor functions that shape attention-related sampling are associated with specific spatiotemporal patterns of spiking activity among specific cell types (Fiebelkorn et al., 2018; Gregoriou et al., 2012). We propose that the spatiotemporal patterns of pre-target spiking activity that occur during trials that result in faster RTs, reflect the recruitment of network nodes and functional units (e.g., functionally defined cell types) associated with the enhancement of sensory processing and the suppression of attentional shifts.

## ACKNOWLEDGMENTS

This work was supported by a training fellowship to I.C.F. (F32EY023465), and by grants from NIMH (R01MH064063, Silvio O. Conte Center (1P50MH109429) and NEI (RO1EY017699, R21EY023565) to S.K.

## AUTHOR CONTRIBUTIONS

Conceptualization, I.C.F. and S.K.; Methodology, I.C.F. and S.K; Investigation, I.C.F.; Formal Analysis, I.C.F.; Resources, S.K.; Funding Acquisition, S.K.; Writing – Original Draft, I.C.F. and S.K.; Writing – Review & Editing, I.C.F. and S.K.

## METHODS

The present study used two male *Macaca fascicularis* monkeys (6–9 years old). The Princeton University Animal Care and Use Committee approved all procedures, which conformed to the National Institutes of Health guidelines for the humane care and use of laboratory animals.

### Behavioral Task

We trained two monkeys to perform a spatial-cueing paradigm. This behavioral task is a variant of the Egly-Driver task (Figure 1A) (Egly et al., 1994). Monkeys self-initiated each trial by pressing and holding down a lever. Following the lever press, a fixation cross (0.5°) appeared at the center of the monitor (eye-monitor distance = 57 cm) and remained there throughout the duration of each trial. After 500–1200 ms, two bar-shaped objects (22° x 4.4°) appeared either oriented horizontally, above and below central fixation, or oriented vertically, to the right and the left of central fixation. The horizontal and vertical orientations were equally likely to occur and the closest edge of each bar was positioned 6.6° from central fixation. After 500–1200 ms, a spatial cue (100 ms) occurred at an end of one of the bar-shaped objects, indicating the location where a subsequent, low-contrast (2.5– 4%) visual target was most likely to occur. On the 78% of trials, following a variable cue-target delay of 300–1600 ms, the visual-target (100 ms) occurred at the cued location. On 12% of trials the visual-target instead occurred at either one of two non-cued locations, which were equidistant from the cued location (i.e., either the non-cued location positioned on the same object, or the non-cued location positioned on the second object). If the monkey detected the target, it received a juice reward for releasing the lever within a window of 150–650 ms. During the remaining trials (10%), no visual-target occurred (i.e., catch trials), and the animals instead received a juice reward for holding down the lever until the screen cleared.

We monitored eye position using an infrared eye tracker (either an Eye-trac 6 at 240 Hz from Applied Science Laboratories or an EyeLink 1000 Plus at 1000 Hz from SR Research), and trials were aborted if eye position deviated by more than one degree from central fixation (i.e., if the monkey broke fixation). Visual stimuli appeared on a 21-inch CRT monitor set at a refresh rate of 100 Hz, and we verified stimulus timing using a customized photodiode system.

### Electrophysiology

We performed all surgical procedures using general anesthesia with isoflurane (induction 2– 5%, maintenance 0.5– 2.5%) and under strictly aseptic conditions. Two customized plastic recording chambers were affixed to head implants for each animal, using titanium skull screws and bone cement. Small craniotomies (4.5 mm diameter) inside these chambers provided access to either frontal or parietal regions. The craniotomies were fit with conical plastic-guide tubes filled with bone wax (Pigarev et al., 2009), which held glass-coated platinum-iridium electrodes (impedance: 5 MΩ) in place between recording sessions. Each recording session spanned a few hours, with up to seven sessions per week. After recordings in the left hemisphere, we moved the recording chambers to the right hemisphere for additional recordings.

During recordings, the animal’s head was stabilized with four thin rods that slid into hollows in the side of the implant. Electrodes were independently lowered with microdrives (NAN Instruments) coupled to an adapter system that allowed different approach angles for each ROI. Electrode signals (40,000 Hz sample rate for spikes; 1,000 Hz sample rate for LFPs) were amplified and filtered (150–8,000 Hz for spikes; 0.7–300 Hz for LFPs) using a Plexon preamplifier with a high input impedance headstage and Multichannel Acquisition Processor (MAP) controlled by RASPUTIN software.

During recordings, we sorted spikes online to isolate neurons, and then resorted for offline analyses using Plexon Offline Sorter software. The present analyses only used neurons with a visual-sensory response (i.e., visual neurons) or both a visual-sensory response and saccade-related activity (i.e., visual-movement neurons). Previous research has established that elevated spike rates during a cue-target delay (i.e., during attentional deployment) are limited to these cell types (Fiebelkorn et al., 2018; Gregoriou et al., 2012; Thompson et al., 2005) (i.e., neurons that demonstrate a visual-sensory response). For within-region analyses, we present data from 82 neurons in FEF and 74 neurons in LIP. For between-region analyses, which required that both brain regions had their strongest responses to stimuli within the same visual-quadrant (i.e., aligned RFs and/or multi-unit RFs), we present data from 82 neurons in FEF and 74 neurons in LIP.

### Acquisition of Structural Images for Electrode Positioning

The monkeys were sedated with ketamine (1-10mg/kg i.m.) and xylazine (1-2 mg/kg i.m.), and provided with atropine (0.04 mg/kg i.m.). Sedation was maintained with tiletamine/zolazepam (1-5mg/kg i.m.). The animals were then placed in an MR-compatible stereotaxic frame (1530M; David Kopf Instruments, Tujunga CA) and vital signs were monitored with wireless ECG and respiration sensors (Siemens AG, Berlin), and a fiber optic temperature probe (FOTS100; Biopac Systems Inc, Goleta CA). Body temperature was maintained with blankets and a warm water re-circulating pump (TP600; Stryker Corp, Kalamazoo MI).

Structural MRI data were collected for the whole brain on a Siemens 3T MAGNETOM Skyra using a Siemens 11-cm loop coil placed above the head. T2-weighted images were acquired with a 3-dimensional turbo spin echo with variable flip-angle echo trains (3D T2-SPACE) sequence (voxel size: 0.5mm, slice orientation: sagittal, slice thickness: 0.5mm, field of view (FoV): 128mm, FoV phase: 79.7%, repetition time (TR): 3390ms, echo time (TE): 386ms, base resolution: 256, acquisition time (TA): 17min 51sec). These images were used both to select coordinates for chamber placements and to position electrodes for recordings. Platinum-iridium electrodes create a clearly identifiable, susceptibility-induced signal void along the length of the electrodes in structural MRI images. This “shadow” has a width of approximately one voxel (0.5 mm^3^ on either side of the electrode), which allows for visualizing electrode placements.

Prior to recordings, electrodes were positioned just above our ROIs. The electrodes were then held *in situ* by customized guide tubes and lowered into cortex over the course of typically one week of recordings. Additional structural MRI data were then acquired prior to replacing the electrodes. Before and after images were used, as well as daily microdrive measurements, to reconstruct electrode tracks.

To further localize electrode penetrations, the D99 digital template atlas was aligned to each individual animal’s MRI volume, using a combination of FSL and AFNI software tools (Cox, 1996; Jenkinson et al., 2012; Reveley et al., 2017). The D99 atlas is based on and aligned to MRI and histological data from the Saleem and Logothetis (2007) atlas, and allows identification of labeled areas within the native 3D MRI volume of an individual animal. Briefly, the brains were first extracted from the MRI volumes using the FSL brain extraction tool (BET) (Smith, 2002). Next, to improve alignment accuracy, the contrast of the MRI volumes was inverted to resemble the image contrast of the T1-weighted atlas MRI volume. The pipeline provided by Reveley et al. (2017) was then implemented to align the atlas to each monkey’s MRI volume. This pipeline included a sequence of affine and nonlinear registration steps to first align the individual animal’s MRI volume to the atlas, then inverting the transformations to warp the atlas to the animal’s original native space. Once aligned, the atlas’ anatomical subdivisions were visually overlaid upon the individual monkey’s MRI volume to assist in the electrode localizations. For all recordings presented here, the electrodes were positioned in atlas-defined FEF and LIP.

## QUANTIFICATION AND STATISTICAL ANALYSIS

See Supplementary Figure 14 in ref (Fiebelkorn et al., 2018) and Supplemental Figure 8 in ref (Fiebelkorn et al., 2019) for demonstrations of common behavioral and electrophysiological effects across the two monkeys. For the present study, as with these previous studies, we combined data from the two animals for all analyses presented in the main text.

### Spike Rate

For all analyses, we used a combination of customized Matlab functions and the Fieldtrip toolbox (Oostenveld et al., 2011) (http://www.ru.nl/neuroimaging/fieldtrip). To estimate changes in spike rate over time, we convolved spikes from each trial with a Gaussian filter (σ = 10 ms) and averaged the resulting functions. To identify visual-sensory neurons, we determined whether there was a statistically significant increase in spike rate in response to the cue (i.e., within 250 ms after cue or target presentation) by using a non-parametric randomization procedure. One response value was randomly selected from the pre-cue period (−350–0 ms) of each trial, averaging those values across trials. This procedure was repeated 5000 times to generate a reference distribution (for the baseline spike rate). The p-value for a non-parametric test is the proportion of values in the reference distribution that exceeds the test statistic (i.e., the observed value from collected data). For all statistical comparisons, unless otherwise specified, we adopted an alpha criterion of 0.05, and used the Holm’s sequential Bonferroni correction to control for multiple comparisons.

To create population PSTHs, we normalized the spike rate for each neuron by its maximum response during trials, and then grand-averaged the normalized spike rates across neurons. To test whether between-condition comparisons were statistically significant, we averaged the response for each neuron over a 500-ms window preceding the target and then used a Wilcoxon rank-sum test. We first used this approach to measure whether spike rate was significantly elevated following the spatial cue and prior to the presentation of the visual target (i.e., during the cue-target delay), comparing spike rate on trials when receptive fields overlapped the cued location relative spike rate on trials when receptive fields overlapped the non-cued location (Figure 1C). We then focused on exploring a potential link between spike rate and response times (RTs), exclusively when receptive fields overlapped the cued location (i.e., under conditions of attentional deployment). That is, we split trials into fast- and slow-RT trials—based on the median RT—and measured the difference in spike rate between these trial types, first combining all visual-sensory neurons (Figure 1D), then separately examining visual and visual-movement cell types (Figure 2). See ref(Fiebelkorn et al., 2018) for the procedure we used to further bin visual-sensory neurons into visual and visual-movement neurons. We also examined potential differences in spike rate for fast- and slow-RT trials by calculating a modulation index for each neuron, subtracting the averaged spike rate on slow-RT trials from the averaged spike rate on fast-RT trials, then dividing by the summed spike rates for each trial type (Figure 1E). We again used a Wilcoxon rank-sum test to determine whether the distribution of values (i.e., the modulation index for each neuron) differed between fast- and slow-RT trials.

### High-frequency Band Activity

To examine a potential link between population spiking and response times, we extracted high-frequency band (HFB) activity from the local field potentials (LFP). Previous research has established that HFB activity is correlated with neuronal firing (Ray and Maunsell, 2011). To isolate HFB activity, we band-pass filtered the raw LFP signal in 8 non-overlapping 10-Hz wide bins from 70–150 Hz. We then calculated amplitude time courses by applying a Hilbert transform to the band-pass filtered data and taking the absolute value. Prior to combining the 10-Hz wide bins, each of the 8 traces was separately baseline corrected by means of a z-score relative to the pre-cue period (i.e., -200–0 ms) (Helfrich et al., 2018). This approach accounts for the 1/f signal drop off as frequency increases, ensuring that HFB activity (70–150 Hz) is not biased toward lower frequencies.

After isolating HFB activity, we first compared trials when the response fields (the LFP-equivalent of receptive fields) overlapped either the cued or the non-cued location. As with the analysis of single-unit activity (i.e., spike rates), we specifically compared HFB activity over a 500-ms window preceding the target using a Wilcoxon rank-sum test to determine statistical significance (Figure 3A). We compared HFB activity on fast-RT trials relative to slow-RT trials, exclusively examining trials when the response fields overlapped with the cued location (i.e., under conditions of attentional deployment (Figure 3B).

### Spike-LFP phase coupling

We used the phase-locking value (PLV) to measure spike-LFP phase coupling. That is, we measured whether spikes were clustering at specific phases of oscillatory activity in LFP, both within a region (i.e., local spike-LFP phase coupling) and between regions (i.e., network-level spike-LFP phase coupling). For each spike time during the cue-target delay (i.e., in a 500-ms window preceding the target), we measured the phase of oscillatory activity in the LFPs across multiple frequencies from 3–55 Hz. To get phase estimates, we convolved the LFP signal with complex wavelets centered on each spike time. We then took the angle of the complex results and measured the consistency of these phase estimates across spike times by calculating PLVs (Lachaux et al., 1999). We first compared PLVs during trials when the receptive fields and response fields overlapped the cued location to PLVs during trials when the receptive fields and response fields overlapped the non-cued location (i.e., attended vs. unattended; Figure 4A). We then compared PLVs for fast-RT trials to PLVs for slow-RT trials, where there was an equal number of trials and near equal number of spikes (Figure 1D) (Vinck et al., 2010), exclusively when the response fields and receptive fields overlapped the cued location (Figure 4–6). As with our analysis of spike rates, we also examined spike-LFP phase coupling separately for visual and visual-movement neurons (Figures 5,6). To visualize spike-LFP phase coupling as a function of time (i.e., at different time points relative to target presentation), we used spikes in 100-ms windows (e.g., from -500 to -400 ms prior to the target), shifting the 100-ms window in 20-ms steps (Figure 7A).

To test for statistically significant between-condition differences in spike-LFP phase coupling, we (i) shuffled (1500 times) phase measurements between conditions (attended vs. unattended trials or fast-RT vs. slow-RT trials), and then (ii) recalculated the difference in PLV between the randomized conditions. We then compared the resulting reference distributions with the observed difference in PLV values.

Between-condition differences in power can lead to spurious differences in spike-LFP phase coupling. We therefore conducted a control analysis that equated power between fast- and slow-RT trials (Supplemental Figure 5). We used the ft_stratify function from the FieldTrip toolbox (Donders Institute for Brain, Cognition, and Behaviour), which also equates sample sizes. Stratification involves subsampling the original dataset to equate power, meaning that the results vary somewhat on each run. We therefore ran 1500 iterations of the stratification procedure at each coupling frequency (from 3–55 Hz, in 1-Hz steps).

Finally, based on previous evidence that higher-frequency activity in the attention network is temporally organized by theta-band activity (Fiebelkorn and Kastner, 2019), we tested whether spike-LFP phase coupling at behaviorally relevant frequencies was modulated by theta phase. Specifically, we examined whether beta-band (19–26 Hz) coupling in FEF was modulated by theta phase and whether alpha-band (9–15 Hz) coupling in LIP was modulated by theta phase. For each spike in the pre-target period, we measured the phase of both higher-frequency activity and theta-band activity. Next we calculated a PLV for the higher frequency band (e.g., the beta band) using a subset of trials spanning a 180° theta-phase window. We then shifted the theta-phase window by 10° and recalculated a PLV for the higher frequency band, based on the new subset of trials. We repeated this procedure until we generated a PLV by theta phase function, spanning all phases. Hypothesizing a signature shape, with a peak in higher-frequency PLV separated from a trough by approximately 180°, we reduced these functions to a single value for each frequency. Specifically, we applied the fast Fourier transform (FFT) to each function (i.e., at each frequency, from 3–60 Hz) and kept the second component, which represents a one-cycle sine wave (matching the hypothesized shape of our PLV by theta phase functions). The amplitude of this one-cycle, sinusoidal component—determined both by how closely the function approximated a one-cycle sine wave and by the effect size—was used to measure the strength of the relationship between theta phase and spike-LFP phase coupling at higher frequencies (Fiebelkorn et al., 2018; Fiebelkorn et al., 2013). To determine statistical significance, we randomly shuffled (1500 times) the theta phase measurements and followed the same procedure to generate a reference distribution.

## SUPPLEMENTAL FIGURES

**Supplemental Figure 1.**
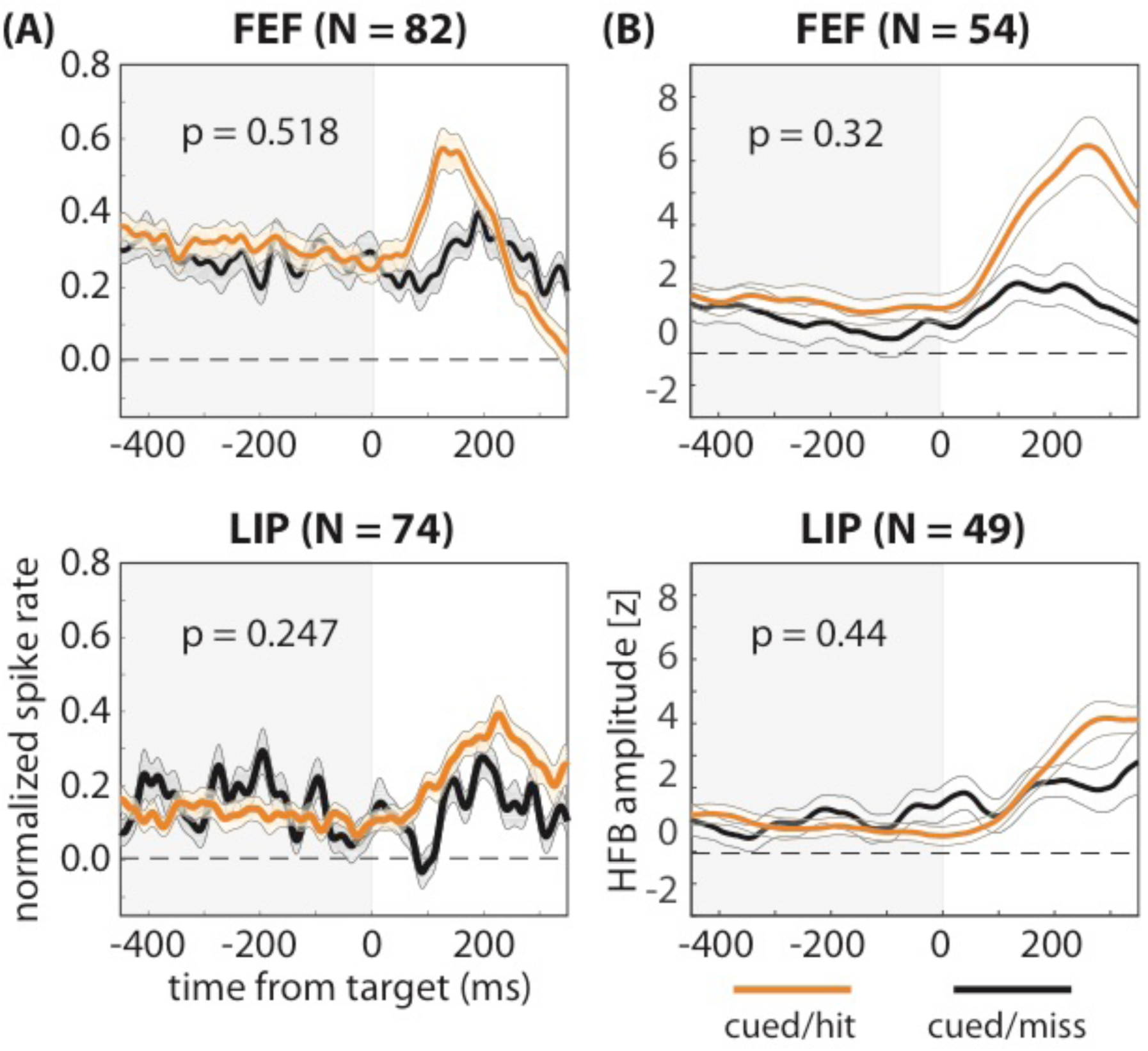
Differences in the magnitude of pre-target spiking activity are not associated with whether a trial resulted in either a hit or a miss. There were no observed differences between trials that resulted in either hits or misses (i.e., when RFs overlapped the cued location) between either **(A)** spike rates averaged across neurons or **(B)** high-frequency band (HFB) activity averaged across sessions (i.e., a proxy for population spiking). The shaded area around each line represents the standard error of the mean.

**Supplemental Figure 2.**
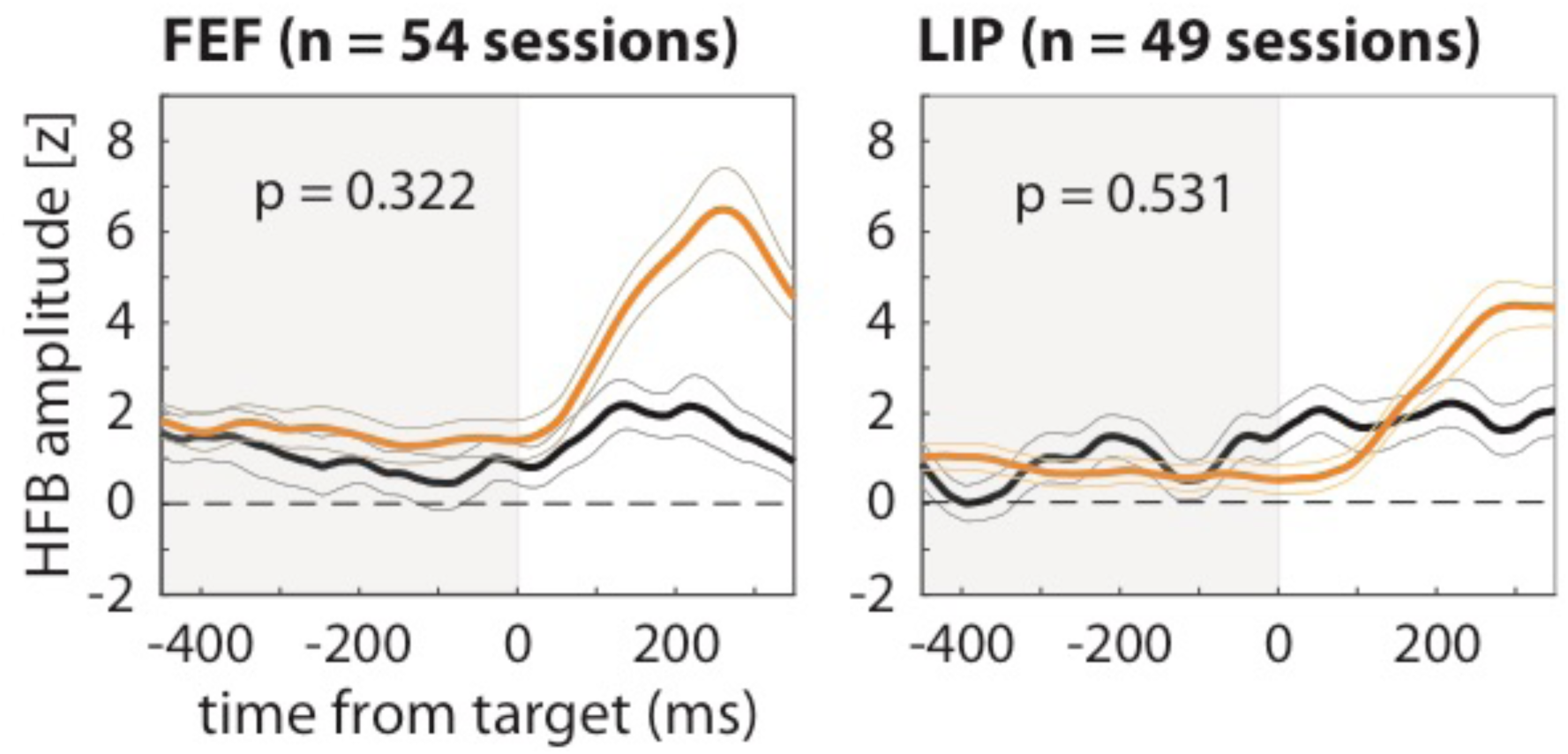
Differences in pre-target population spiking are not associated with differences in behavioral performance. We observed no differences in HFB activity between trials that resulted in either hits (orange lines) or misses (black lines). The shaded area around each line represents the standard error of the mean.

**Supplemental Figure 3.**
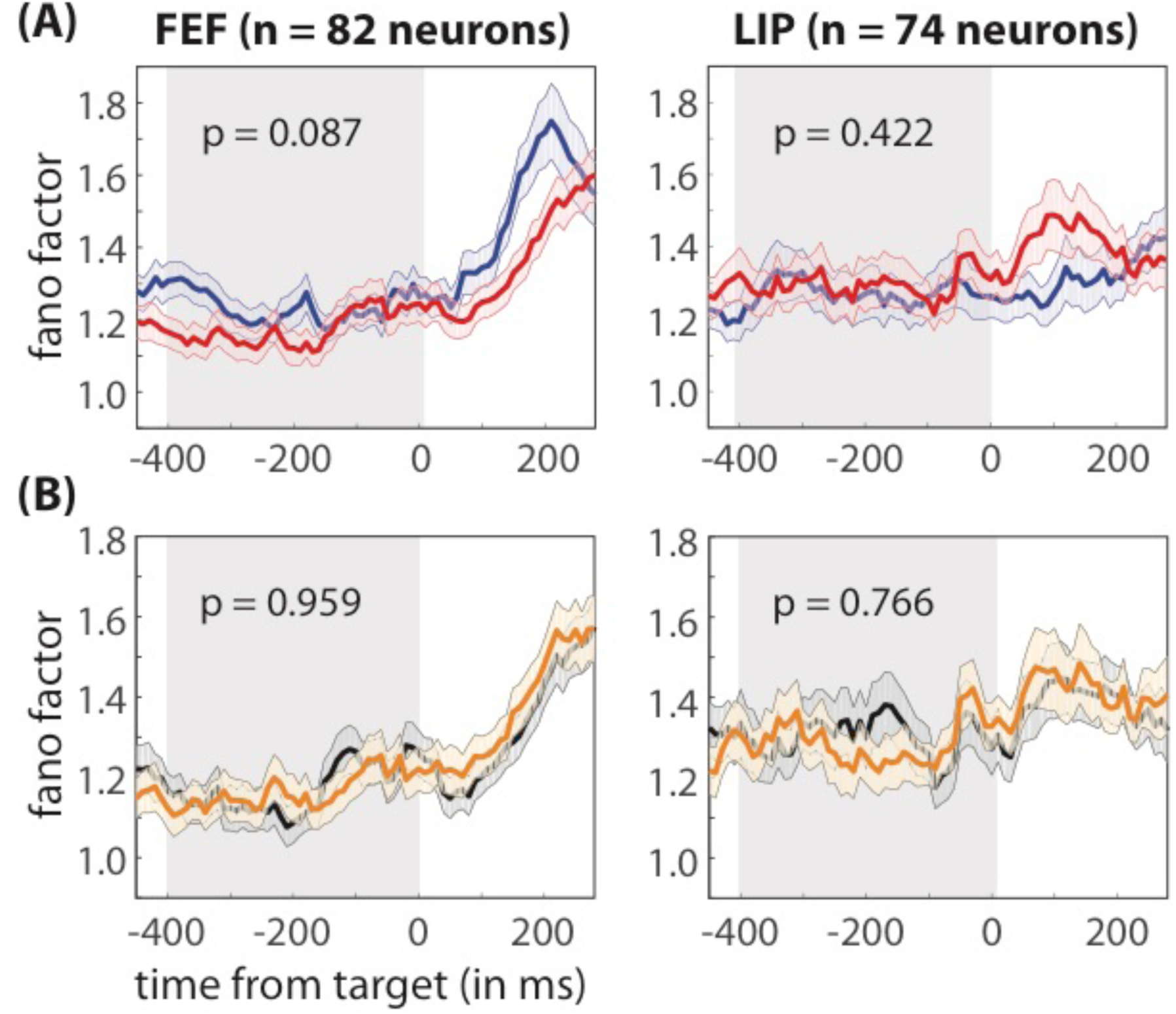
Differences in pre-target spike-count variability are not associated with differences in behavioral performances. There was no difference in spike count variability (i.e., Fano factor) based on either **(A)** attentional condition (i.e., cued versus non-cued) or **(B)** behavioral performance (i.e., fast versus slow RTs). The shaded area around each line represents the standard error of the mean.

**Supplemental Figure 4.**
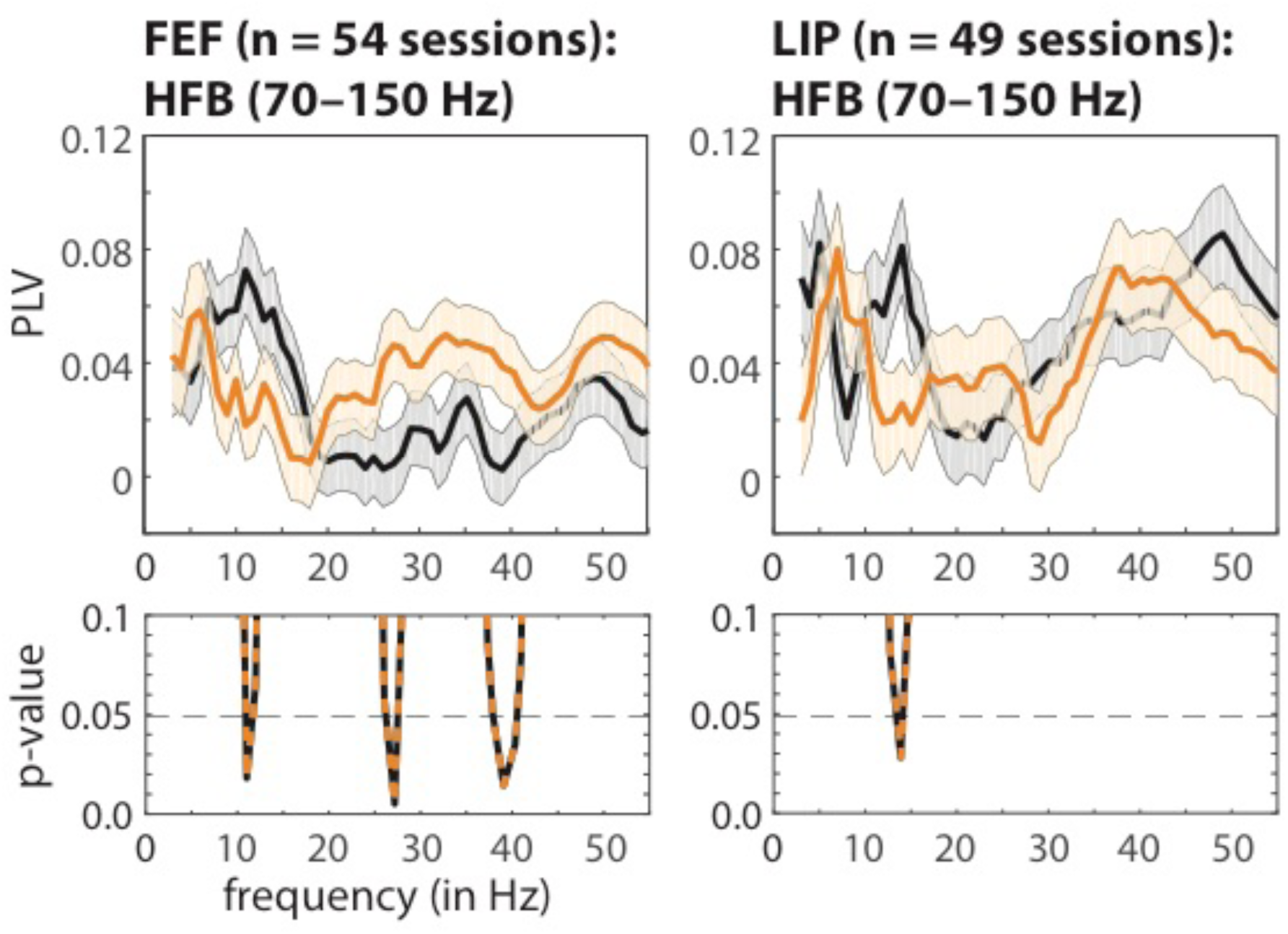
Different oscillatory patterns in pre-target population spiking activity are associated with either faster or slower response times. We compared the consistency of oscillatory phase at peaks in high-frequency band (HFB) activity, during the cue-target delay, across fast- (orange lines) and slow-RT trials (black lines). These results are comparable to those associated with local spike-LFP phase coupling (Figure 4). The p-values for between-condition comparisons are represented below each panel. The shaded area around each line represents the standard error of the mean.

**Supplemental Figure 5.**
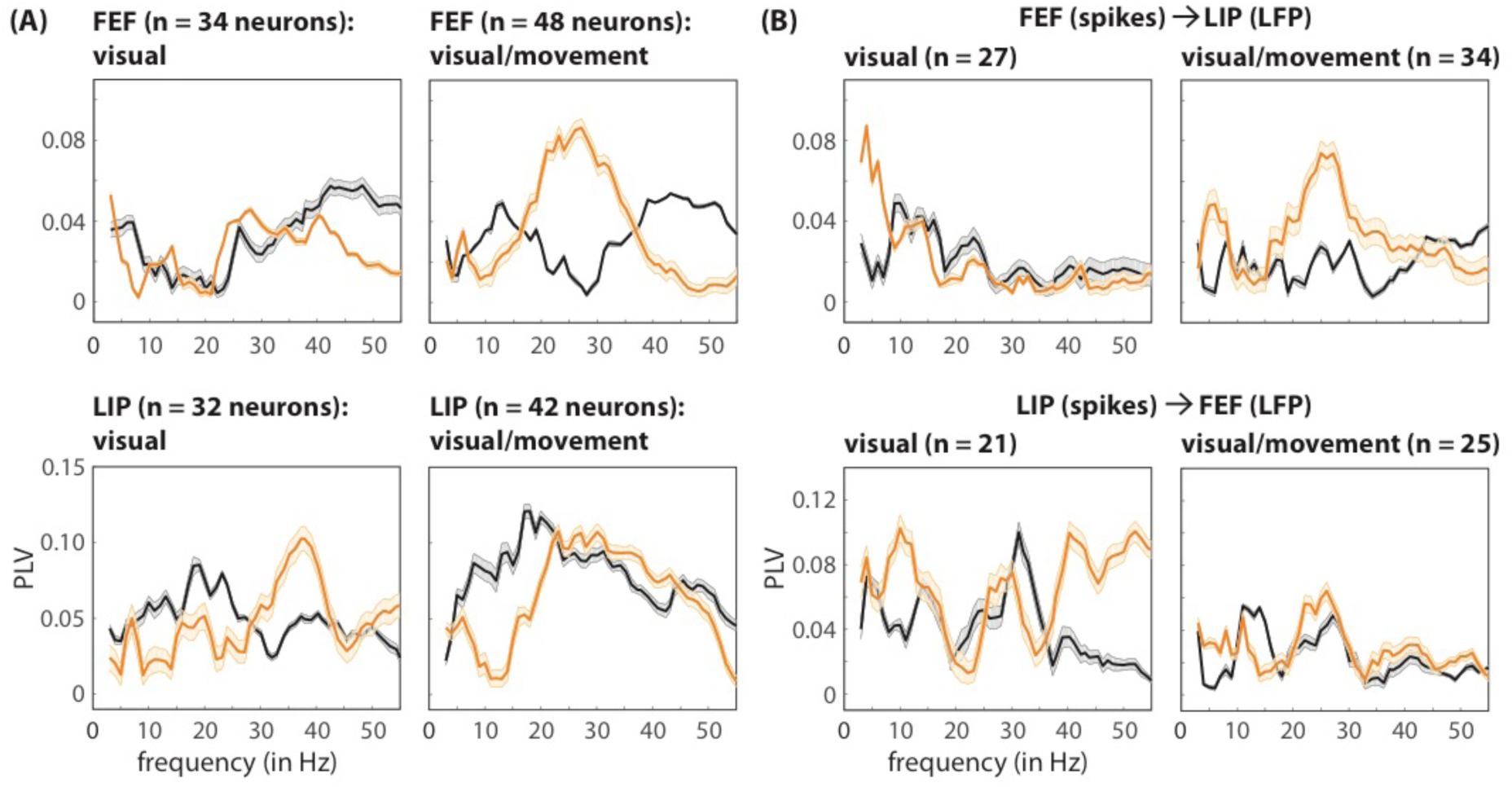
Behaviorally relevant differences in oscillatory patterns of spiking activity are not attributable to differences in oscillatory power. The reliability of phase estimates increases with increasing oscillatory power. This may lead, for example, to spurious between-condition differences in spike-LFP phase coupling if there are between-condition differences in oscillatory power. Here, differences in spike-LFP phase coupling, at both the **(A)** local and the **(B)** network levels, continued to be observed after a stratification procedure that equated power across trials that resulted in either faster or slower response times. The shaded area around each line represents the standard error of the mean associated with repeated iterations of the stratification procedure.

